# The dentate gyrus grows throughout life despite turnover of developmentally-born neurons

**DOI:** 10.64898/2026.02.18.706677

**Authors:** Tina Ciric, Shaina P Cahill, Tyler Lin, Si-ah Choi, Jason S Snyder

## Abstract

Adult-born hippocampal neurons are highly plastic but there remains uncertainty about the magnitude of neurogenesis and its long-term functional consequences. Theoretical predictions indicate that adult neurogenesis should lead to substantial growth of the dentate gyrus (DG) granule cell population. However, in practice, most studies find no changes in total cell number across adulthood. This discrepancy may partly be a sensitivity issue, where small sample sizes and the examination of older age windows (when neurogenesis is reduced) have prevented detection. However, neurogenic growth could also be masked by the turnover of developmentally-born DG neurons, which are known to die off in normal aging. To address the question of how neuronal birth and loss impacts DG population dynamics, here we quantified numbers of developmentally-born neurons, proliferating Ki67+ cells (as a proxy for adult-born neurons), and total DG neurons from 2-18 months of age in the rat. We estimate that over this timeframe 670,000 adult-born neurons are added (30% of the total population). Consistent with neurogenic growth, the total number of DG neurons increased across adulthood. However, net growth was only 385,000 cells, which is less than predicted by adult neurogenesis alone. Indeed, 20% of developmentally-born neurons were lost over the same interval, and so we propose that the difference is explained by neuronal turnover. Neuronal persistence and turnover may be relevant for theories of hippocampal long-term memory, as well as for understanding psychiatric conditions that are characterized by hippocampal plasticity and atrophy.

## Introduction

The magnitude and significance of adult hippocampal neurogenesis has been debated since its discovery by Joseph Altman in the 1960s (Altman and Das, 1965; Gross, 2000; Snyder, 2019). While its occurrence in animals isn’t contested, even in rodents it is often stated that adult neurogenesis produces only a small number of dentate gyrus (DG) neurons. This perception may be because histological markers and transgenic mice typically only label a snapshot of neurons born in a short temporal window. When short-term neuronal production is integrated over time, simple math predicts that the cumulative effects of adult neurogenesis could end up producing as much as 40-50% of DG neurons over the lifespan (Snyder and Cameron, 2012; Cole et al., 2020). Indeed, data from transgenic mice that more effectively label stem cell progeny is more consistent with substantial cumulative neurogenesis in adulthood (DeCarolis et al., 2013).

If adult neurogenesis produces a large number of DG granule neurons, one would expect the total number of neurons to increase over adulthood. Seminal work by Bayer provided the first evidence for DG neuron accumulation, but this was over an interval that included the juvenile period (Bayer et al., 1982). Moreover, subsequent work over the same interval (Boss et al., 1985), or exclusively in adulthood (Rapp and Gallagher, 1996; Rasmussen et al., 1996; Kempermann et al., 1998; Merrill et al., 2003), failed to observe DG growth. However, these studies were not optimized to detect DG neuron accumulation: either they did not use efficient and unbiased stereological counting principles, they were underpowered, or they only examined growth at older ages when neurogenesis had declined (indeed, some were designed to detect age-related cell loss rather than cell addition).

Another challenge for detecting cell accumulation is the fact that mature, developmentally-born neurons (DBNs) die off in adulthood as a part of normal aging (Dayer et al., 2003; Cahill et al., 2017; Ciric et al., 2019). Thus, when counting total DG neurons, net growth due to adult neurogenesis may be masked by neuronal loss. This possibility is supported by another well-established model of neurogenesis, the avian HVC-RA song circuit, where there are seasonal increases in both neurogenesis and cell death (Kirn and Nottebohm, 1993). Functionally, this interrelated process of neuronal death and addition supports the degeneration and regeneration of song (Cohen et al., 2016). And, while such seasonal/stereotyped patterns of memory turnover are less apparent in mammals, it may be that some memories are cleared from the hippocampus but others find permanent storage there (Moscovitch and Gilboa, 2021). However, it is unclear whether there exist patterns of neuron addition and turnover that could support different memory dynamics.

To address these gaps, here we characterized the dynamics of DG neuron addition and loss in a large rat cohort. We quantified the loss of DBNs, the addition of new adult-born DG neurons (ABNs), and the total number of DG across young adulthood through old age. Our findings indicate that, while a substantial number of neurons turn over, cumulative neurogenesis results in net growth of the DG granule cell population in adulthood.

## Methods

### Animals and treatments

All animal procedures adhered to Canadian Council on Animal Care guidelines and were approved by the University of British Columbia Animal Care Committee. Long-Evans rats were bred and housed in the Department of Psychology animal facility under a 12:12 light/dark cycle (lights on at 6:00 a.m.). Breeding pairs were maintained in large polyurethane cages (47 × 37 × 21 cm) containing a polycarbonate shelter tube, aspen bedding, and unrestricted access to chow and water. Male breeders were removed from the cage prior to birth. The day of birth was designated postnatal day (P) 1 and pups weaned at P21, at which point they were separated into pairs and transferred to smaller polyurethane cages (48 × 27 × 20 cm). All experimental animals were born in May-June 2015. Only male rats were used for experiments, as was typical at the time (Shansky and Murphy, 2021; Knudson et al., 2022).

On P6, all rats received an injection of CldU (50mg/kg, intraperitoneal). Rats were then allowed to survive to 2, 4, 6, 12 or 18 months of age to track the long-term survival of developmentally-born cells (Fig. 1). Four weeks prior to endpoint, rats were injected with IdU (100mg/kg, intraperitoneal), to quantify adult neurogenesis with a different S-phase marker in the same animals (Cahill et al., 2018). However, we could not obtain IdU-specific immunostaining and so these cells were not analyzed as a part of this project. Offspring from each litter were distributed evenly across experimental groups.

**Figure 1:**
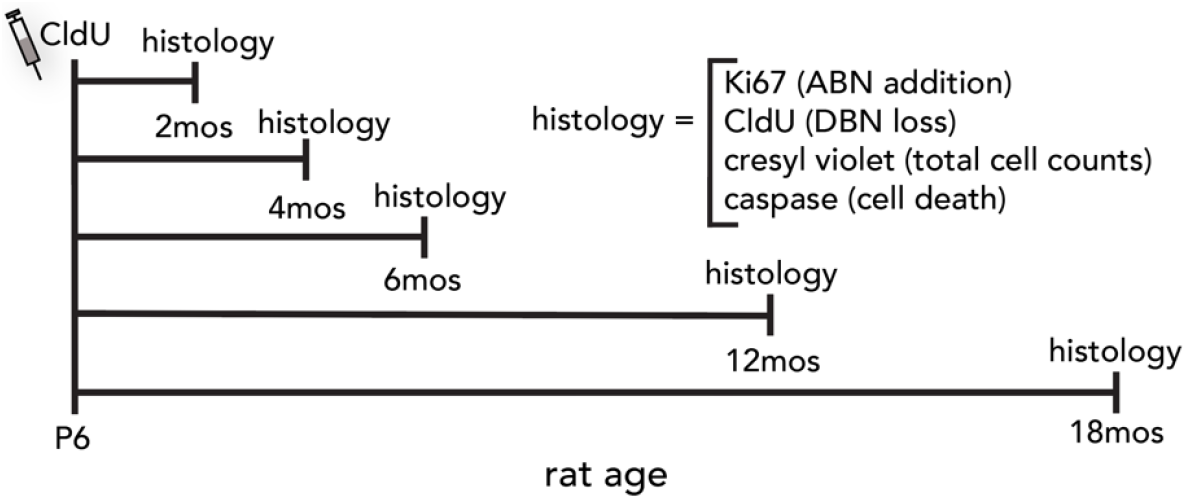
Experimental design. Rats were injected with the mitotic label, CldU, on postnatal day 6 (P6). At 2,4,6,12 or 18 months of age, brains were collected and stained for the proliferation marker Ki67 (to quantify adult-born neurons; ABNs), CldU (to quantify surviving developmentally-born neurons; DBNs), or cresyl violet (to quantify total numbers of dentate gyrus granule cells).

### Tissue preparation and immunohistochemistry

Rats were transcardially perfused with 4% paraformaldehyde in phosphate-buffered saline (PBS; pH 7.4). Brains were post-fixed in PFA for an additional 48 hours and then transferred to PBS containing 0.1% sodium azide for storage prior to histological processing. For cryoprotection, brains were sequentially immersed in 10% glycerol (24 hr) followed by 20% glycerol (48 hr). Coronal sections (40 µm) were cut on a freezing microtome and stored at –20°C in cryoprotectant solution until immunostaining.

For Ki67 cell counting, a 1-in-12 series of sections spanning the full rostrocaudal extent of the dentate gyrus was mounted on slides. Slides were heated to 90°C for 10 minutes in citric acid (0.1M, pH 6.0) for antigen retrieval, washed in PBS, and then incubated with mouse anti-Ki67 antibody for 24 hours (1:200 in PBS containing 0.5% triton-x and 3% horse serum; BD Biosciences #550609). Tissue was then washed with PBS and incubated with biotinylated goat anti-mouse secondary antibody (1:250; Sigma #B0529) for 1 hour. Ki67^+^ cells were visualized using an avidin–biotin HRP detection kit (Vector Laboratories) and cobalt-enhanced DAB (Sigma Fast Tablets). Slides were rinsed in PBS, dehydrated through graded ethanol solutions, cleared in Citrisolv (Fisher), and coverslipped with Permount.

For CldU cell counting, a 1-in-12 series of sections spanning the full rostrocaudal extent of the dentate gyrus was mounted on slides. Slides were heated in 0.1 M citric acid (pH 6.0) at 90°C, treated with trypsin, and incubated in 2N HCl for 30 minutes to denature DNA. Sections were then incubated overnight with rat anti-BrdU (1:1000; Serotec # OBT0030G) diluted in PBS containing 10% triton-x and 3% horse serum. The following day, tissue was washed and incubated with biotinylated donkey anti-rat secondary antibody (1:250; Jackson Immuno 712-065-153) for 1 hour. CldU^+^ nuclei were visualized using an avidin–biotin HRP detection kit (Vector Laboratories) and cobalt-enhanced DAB (Sigma Fast Tablets). Slides were rinsed in PBS, dehydrated through graded ethanol solutions, cleared in Citrisolv (Fisher), and coverslipped with Permount. Since our experimental rats were also injected with IdU, we stained tissue from rats that received IdU injections but not CldU injections. These control tests revealed no labelled cells and therefore confirm that our CldU staining specifically labelled CldU^+^ cells and not IdU^+^ cells.

For cresyl violet staining, a 1 in 12 series of sections, spanning the full rostrocaudal extent of the dentate gyrus, was mounted onto slides. Slides were briefly rinsed in distilled water and then placed in 0.1% cresyl violet acetate solution for 5 minutes. Slides were then rinsed in distilled water, placed in 70% ethanol for 1 minute, and incubated in acetic acid (0.25% in 95% ethanol) to remove nonspecific cresyl violet signal. Slides were then dehydrated in 100% ethanol, cleared with Citrasolv (Fisher), and coverslipped with Permount.

For caspase-3 staining, free-floating sections were incubated in 2N HCl for 30min, washed, and incubated for 72 hours at 4°C in PBS containing 0.5% triton-x, 3% horse serum, and rabbit anti–active caspase-3 (1:200; BD Biosciences #559565) and rat anti-BrdU antibodies. After washes, sections were incubated for 1 hour at room temperature with Alexa555-conjugated donkey anti-rabbit secondary antibody and Alexa488-conjugated donkey anti-rat secondary antibody (1:250; ThermoFisher). Tissue was counterstained with DAPI, mounted, and coverslipped with PVA-DABCO.

### Microscopy and quantification

CldU-labeled cells were quantified under brightfield illumination using stereological principles. A 1-in-12 series of sections encompassing the entire dentate gyrus was examined with a 40x objective on an Olympus CX41 microscope. All CldU^+^ cells within the granule cell layer were counted. Total CldU^+^ cell number per dentate gyrus (bilateral) was estimated by multiplying the counted values by 12.

Ki67^+^ cells were quantified in the same manner as CldU^+^ cells with the exception that only cells located in the SGZ and deepest aspect of the granule cell layer (i.e. where it borders the hilus) were quantified.

Cresyl violet-stained granule cells were quantified using stereological principles and the optical fractionator method (West et al., 1991). An automated stage-driven Olympus BX51 microscope and Stereoinvestigator software (MBF Bioscience) was used to systematically and efficiently sample cells throughout the DG and estimate the total number of granule cells according to the equation: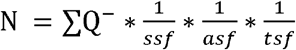, where ∑Q^-^is the total number of cells counted per rat, ssf is the section sampling fraction, asf is the area sampling fraction, and tsf is the tissue thickness sampling fraction. For each rat, a 1 in 12 series of sections was examined (thus, the SSF was 1/12), the counting frame was 11.8 × 11.8 µm, the sampling grid was 250 × 250 µm (thus the asf was 0.0022), and the dissector height was 4 µm with a guard zone of 2 µm. The tissue thickness sampling fraction was determined by the ratio of the dissector height to the local number-weighted mean section thickness. An average of 171 cells across an average of 153 sampling sites were counted per rat. Optical fractionator efficacy was assessed by computing the sampling/measurement variance (squared coefficient of error, CE^2^) and the biological group variance (squared coefficient of variation, CV^2^). CE (Gundersen, m=1) was 0.08 (CE^2^ = 0.006) for each group, and CV^2^ ranged from 0.02 to 0.05. Thus, our sampling variance was minor in comparison to the age-related group variance, indicating that our sampling scheme was efficient (Slomianka and West, 2005).

Fluorescent caspase-3–stained tissue was imaged with a Leica SP8 confocal microscope. For analyses of apoptotic cells in 2-, 12-, and 18-month-old animals, only cells that showed both strong caspase-3 immunoreactivity and pyknotic morphology were included. A 1-in-12 series of sections was screened for caspase^+^ cells and for caspase^+^CldU^+^ cells. Cell counts were normalized to the 2-dimensional area of the granule cell layer and expressed as a density measurement.

To infer the approximate age of dying granule neurons, the position of each pyknotic caspase^+^ cell was mapped within the granule cell layer. Consistent with the known inside-out gradient of granule cell maturation (Crespo et al., 1986; Wang et al., 2000; Mathews et al., 2010; Cahill et al., 2017), immature neurons were expected to lie near the hilus/subgranular zone and older cells near the molecular layer. For each section, granule cell layer thickness was normalized to a 0–100 scale, where 0 represented the hilar border and 100 the molecular layer border. The position of each dying cell was recorded as its proportional distance from the hilar boundary. Cells located in the subgranular zone were assigned a value of 0.

### Statistical analyses

Data were analysed with Prism (Graphpad) and R. Ki67 cell counts were analyzed by 1-way ANOVA followed by Holm-corrected multiple comparison tests between all ages.

Ki67, CldU and cresyl violet cell counts were fit with linear, quadratic and cubic polynomial equations and the best fitting model that was also statistically better than simpler fits was chosen to estimate cell addition/loss (Ki67: cubic, CldU and cresyl violet: linear). Caspase^+^ cell densities were analyzed by 2-way ANOVA (region x age) and caspase^+^ cell locations in the granule cell layer were analyzed by 1-way ANOVA, with Holm-corrected multiple comparisons tests. For all tests, ⍰ = 0.05.

## Results

To estimate the amount of neurogenesis throughout the lifespan we counted total proliferating Ki67^+^ cells present in the DG of 2,4,6,12, and 18 month-old rats (Fig. 2). As expected, there was an exponential decline in Ki67^+^ cells that plateaued by 12 months of age (ANOVA, effect of age: F_4,64_=49, P<0.0001; all ages significantly different from each other (P<0.01) except 12 months vs 18 months (P=0.6)). The age-related changes in proliferation were best fit by a cubic polynomial equation (R^2^ = 0.75), which we used to estimate that 670,000 cells were added from 2 to 18 months of age (assuming each Ki67^+^ cell translates to a surviving ABN).

**Figure 2:**
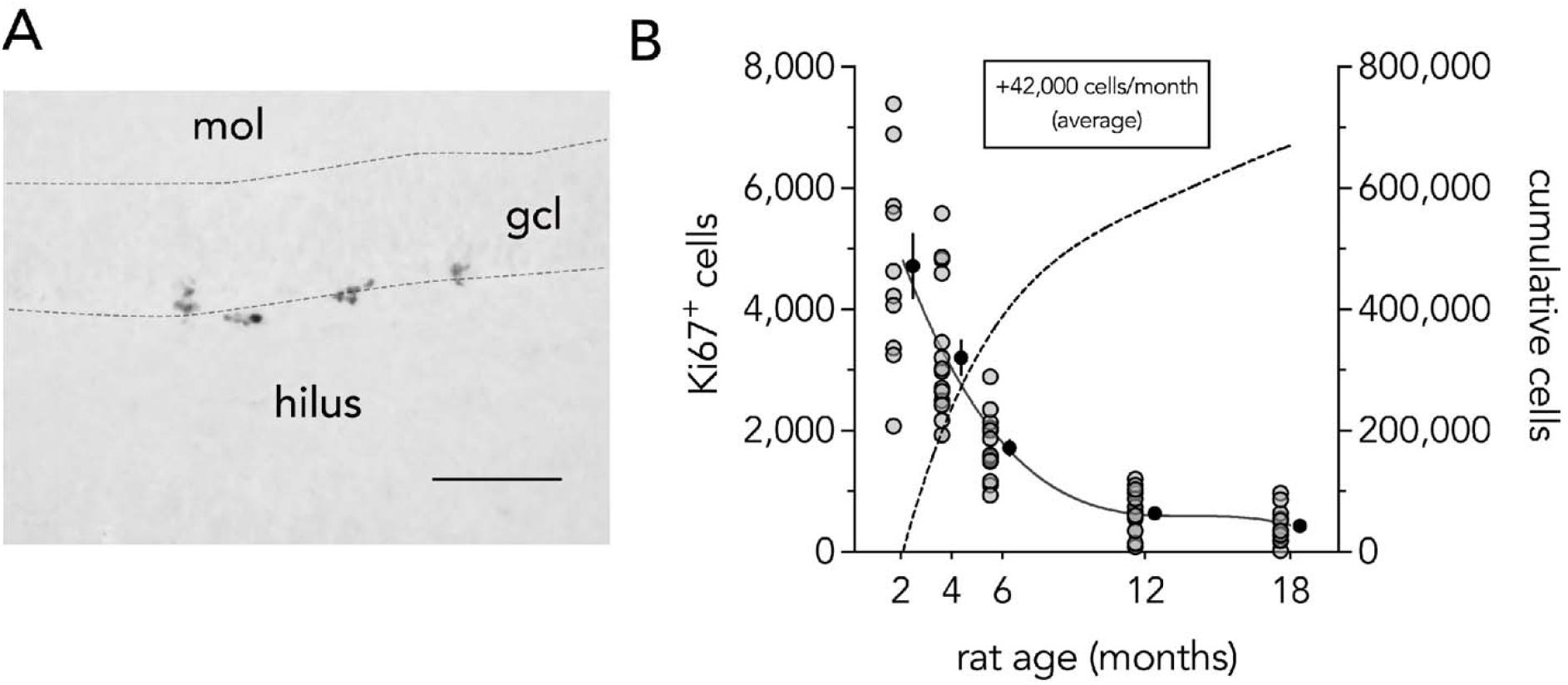
Neurogenesis across the lifespan. Left: clusters of proliferating Ki67^+^ cells at the border of the granule cell layer and the hilus (subgranular zone) of the dentate gyrus. Scale bar = 100 µm. Right: Ki67^+^ cells decline with age and plateau by 12 months. Solid black line reflects cubic fit of the data. Grey symbols indicate cell counts for individual rats, black symbols indicate group means ± standard error. Dashed line is the integral of the cubic fit, to estimate the total number of adult-born cells added over the lifespan (right y-axis). mol, molecular layer; gcl, granule cell layer.

We and others have previously observed death of P6-born DBNs that are born at the peak of rat DG development (Dayer et al., 2003; Cahill et al., 2017; Ciric et al., 2019). To examine whether these cells continue to die throughout the rest of the lifespan, they were labelled with CldU and counted at 2,4,6,12 and 18 months of age (Fig. 3). CldU^+^ cells were located throughout the granule cell layer, with a large fraction away from the germinal SGZ and in the superficial portions of the granule cell layer, reflecting their developmental origin. Linear regression revealed a significant age-related decline in the number of cells: ∼1900 P6-born cells lost/month, for a total of ∼30,000 cells over 2 to 18 months of age (19% loss; R^2^=0.09, P=0.006).

**Figure 3:**
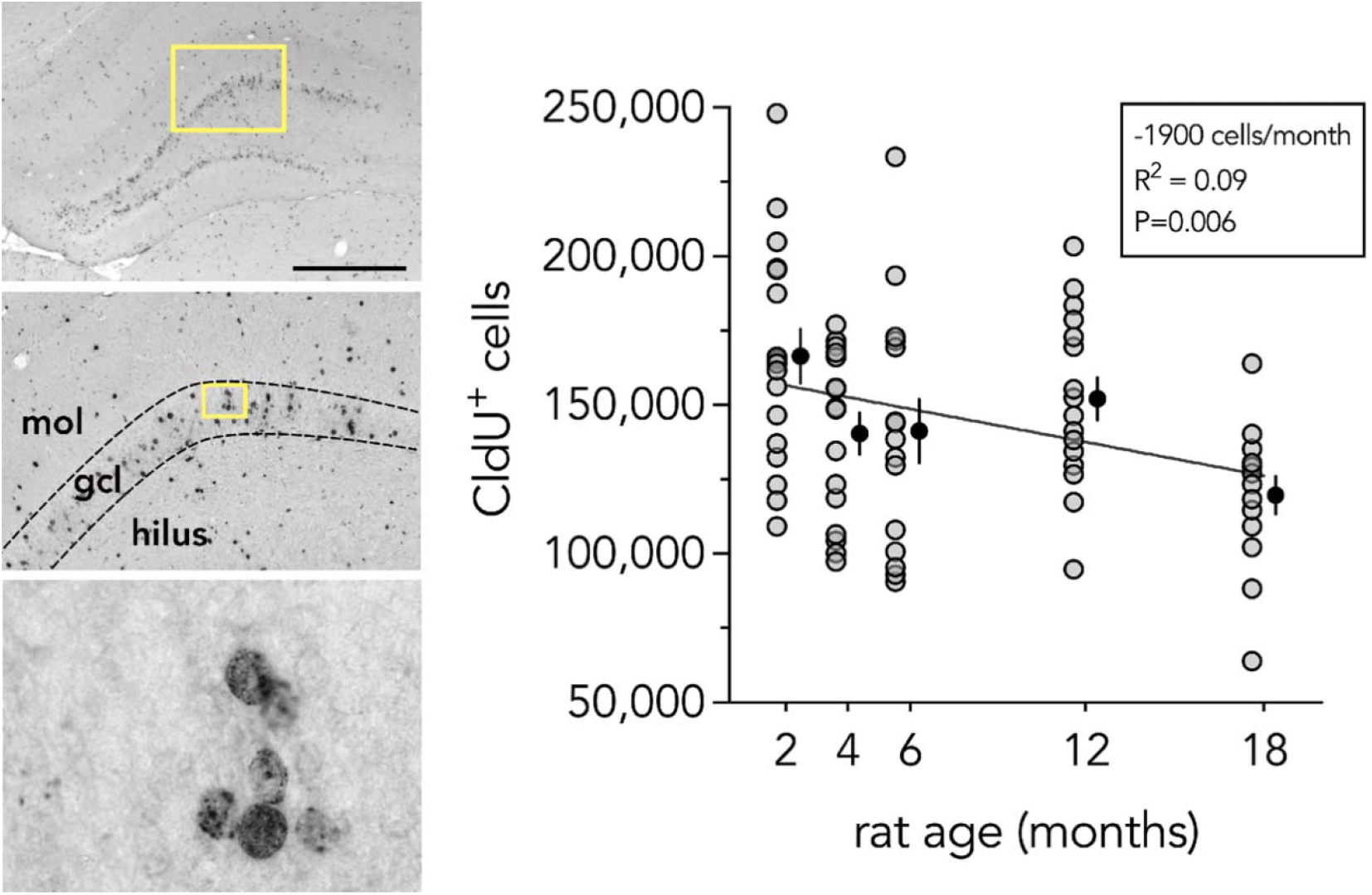
Loss of developmentally-born neurons in aging. Left: images of a rat dentate gyrus immunostained for developmentally-born CldU^+^ neurons. Scale bar = 1000 µm. Yellow boxes indicate regions that are further magnified. Right: CldU^+^ neurons, born on postnatal day 6, are lost throughout adulthood. Line reflects linear regression, which estimates 30,000 cells are lost from 2 to 18 months. Grey symbols indicate individual rats, black symbols indicate mean ± standard error. mol, molecular layer; gcl, granule cell layer.

We next stained for the apoptosis marker, activated caspase-3, to obtain direct evidence for cell death. Focussing on the 2, 12, and 18 month-old groups we counted cells that expressed caspase and were also pyknotic, as reflected by condensed DAPI^+^ chromatin (Fig. 4). Due to the rapid clearance of apoptotic cells, dying cells were rare and there were too few CldU^+^caspase^+^ cells to conduct a proper analysis of birthdated DBNs. However, dying cells were present in both the SGZ (where neurogenesis occurs) and the GCL (where granule neurons reside). Analyses revealed significantly more cell death in both DG regions at 2 months of age than at 12 and 18 months, with comparable rates of cell death across these 2 regions (effect of region, F_1,27_=0.4, P=0.5; effect of age, F_2,27_=53, P<0.0001; region x age interaction, F_1,27_=1.6, P=0.2; 2 months vs 12/18 months both P<0.0001, 12 vs 18 months P=0.4).

**Figure 4:**
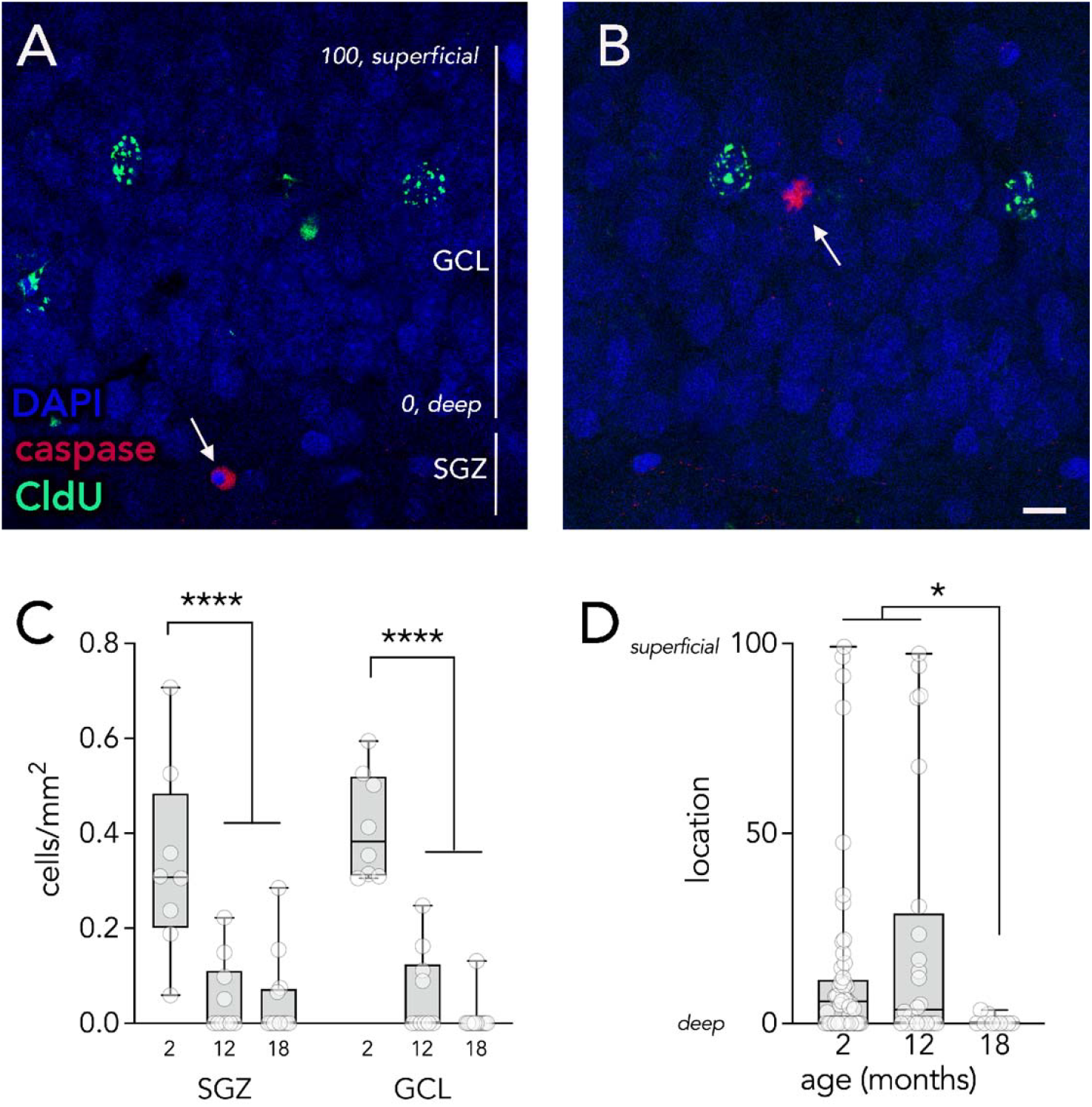
Apoptosis in the dentate gyrus. A) A caspase^+^ cell in the SGZ (arrow), likely reflecting a dying immature neuron. Vertical bars indicate the SGZ and deep-to-superficial range of the GCL. B) A caspase^+^ cell in the superficial granule cell layer (arrow), likely reflecting a dying developmentally-born neuron. SGZ = 0, superficial molecular layer border of the GCL = 100. Scale bar = 10 µm. C) There were more caspase^+^ cells in 2 month old rats than in 12 and 18 month old rats, in both the SGZ and GCL. D) In 2 and 12 month old rats, caspase^+^ cells were located in more superficial layers of the GCL, as compared to 18 month old rats where dying cells were almost exclusively observed in the SGZ. Boxes and whiskers reflect quartiles, symbols indicate individual cells. SGZ, subgranular zone. GCL, granule cell layer. *P<0.05, ****P<0.0001.

**Figure 5:**
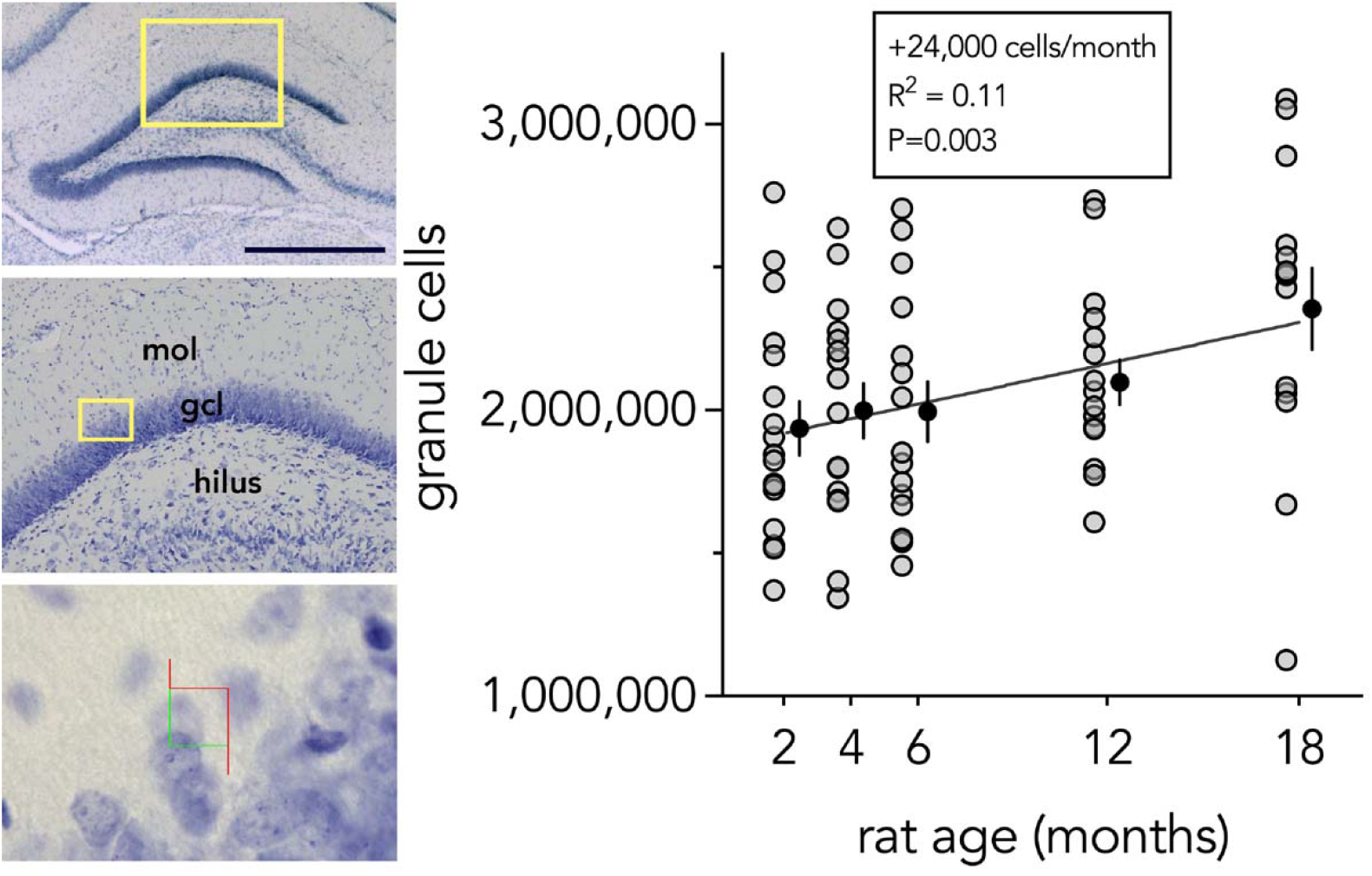
Net addition of granule cells in adulthood. Left: Images of the DG granule cell layer stained for cresyl violet. Bottom image shows the stereological counting frame. Scale bar = 1000 µm. Right: Linear regression estimates net growth of 385,000 granule cells from 2-18 months of age. Grey symbols indicate individual rats, black symbols indicate mean ± standard error. mol, molecular layer; gcl, granule cell layer.

Dentate granule cells are organized by age, where the most immature granule cells are located in the deep portions of the granule cell layer (near the SGZ) and older cells are located in more superficial layers (Crespo et al., 1986). We can therefore obtain reasonable evidence for/against the death of DBNs based on the location of dying cells. We therefore measured the location of pyknotic caspase^+^ cells with respect to the width of the granule cell layer and expressed cell position as a percentage, where 0 reflects a location in the SGZ or deepest cell layer of the GCL and 100 reflects positioning in the most superficial layer of the GCL, at the border of the molecular layer. The majority of cells were located in the SGZ or deep (25%) granule cell layer, consistent with the death of immature ABNs that are dying during their critical period of cell death (0-4w of cell age)(Cameron et al., 1993; Tashiro et al., 2006). However, a portion of cells in 2 and 12 month old rats were located more superficially in the GCL (the superficial 75%), which does not contain immature DCX^+^ cells (Cahill et al., 2017) and therefore is likely to reflect the death of older, mature granule neurons, including DBNs. Overall, 2 and 12 month old rats tended to possess dying cells in more superficial aspects of the granule cell layer as compared to 18 month old rats (Kruskal-Wallis H=7.7, P=0.02; 2 and 12 months vs 18 months both P<0.03, 2 months vs 12 months P=1).

The addition of a large number of cells in adulthood, coupled with the loss of cells born in development, raises the question of whether the total granule cell population grows, shrinks or remains constant with age. To address this question, we stained the DG for cresyl violet and counted total numbers of granule neurons using the optical fractionator method (West et al., 1991). A linear model revealed a strong effect of age, estimating that ∼24,000 cells are added per month, resulting in total growth of 385,000 cells from 2 to 18 months (a 20% increase; R^2^=0.11, P=0.003).

## Discussion

Animal models have provided little direct evidence that neurogenesis makes large-scale contributions to hippocampal structure. By drawing from a large rat sample, and quantifying markers of ABN addition, DBN loss and total granule cell counts, here we show that there is considerable growth of the granule cell population from 2 to 18 months of age, despite the turnover of cells born at the peak of DG development.

Specifically, the total rat granule cell population grows by 385,000 granule cells (a 20% increase), which is substantially less than the ∼670,000 ABNs that are added over the same interval. We propose that the difference, approximately 285,000 cells, reflects the death of DBNs that are born over ∼10 days during the peak of DG development (since 30,000 DBNs, born on a single day, were lost).

### Caveats and assumptions

Technical limitations make it impossible to track the full addition and loss of DG neurons throughout the lifespan. Here, we estimated cell loss and addition based on histological “snapshots” across subjects, and so it is worth addressing assumptions and potential biases that could have influenced our calculations. With respect to total DG cell counts, the cresyl violet cell data are perhaps the clearest evidence that there is neurogenesis-dependent accumulation of DG granule neurons throughout the lifespan. Cresyl violet stains all granule neurons, and so there are no errors introduced by estimating how the histologically stained sample relates to the full population (in contrast to neurogenesis markers such as BrdU, which only label a fraction of ABNs).

While it is possible that systematic biases might have resulted in under/over estimation, this would equally affect all groups. Thus, if nothing else, our data convincingly show that the DG grows throughout life due to adult neurogenesis.

We used the proliferation marker, Ki67, to estimate neurogenesis rates throughout the lifespan (Kee et al., 2002). We assumed that each Ki67^+^ cell resulted in one new granule neuron added to the DG. Despite the complex and multistep nature of neurogenesis (Kempermann et al., 2004), our Ki67^+^ counts are very similar to our previous counts of long-term surviving BrdU^+^ cells (2-month-old rats: ∼5000 Ki67^+^ cells vs ∼6000 BrdU^+^ cells)(Snyder et al., 2009). Since those BrdU cell counts reflected cells labelled by a single injection, which would only label approximately half of the cells born on a given day (Cameron and McKay, 2001), our Ki67 counts may underestimate the true magnitude of neurogenesis. On the other hand, Ki67 may overestimate neurogenesis in aging, when neuronal differentiation rates decline (Gil-Mohapel et al., 2013). For these reasons, we have not attempted to further correct our neurogenesis estimates but treat them as a faithful, if rough, estimate that is consistent with previous models (Snyder and Cameron, 2012; Cole et al., 2020) and empirical evidence (Cameron and McKay, 2001; DeCarolis et al., 2013) that adult neurogenesis makes a substantial contribution to the DG neuronal population.

Finally, we used the thymidine analog, CldU, to label DG neurons born at the peak of DG development, and quantify their loss throughout life. A single CldU injection is unlikely to capture all cells born on a single day (Cameron and McKay, 2001), but it will also label cells born on subsequent days as stem cells redivide (Dayer et al., 2003). Thus, our CldU^+^ cell counts are likely to approximate the number of cells born on a single day. The decline in CldU^+^ cells is unlikely to reflect loss of label (e.g. due to redivision of precursor cells and label dilution (Mathews et al., 2010)) since we have previously observed mature birthdated DBNs, located outside of the neurogenic niche, actively undergoing apoptosis (Cahill et al., 2017). Thymidine analog-induced toxicity, as proposed for the olfactory bulb (Platel et al., 2019), is also unlikely to be a confound for our analyses of DG neuron survival for several reasons. First, immature DG neurons display a critical period of cell survival independently of thymidine analog labelling (Tashiro et al., 2006). Second, BrdU-labelled ABNs and DBNs that are born at other stages of development do not die after cells reach 1 month of age (Dayer et al., 2003; Kempermann et al., 2003; Ciric et al., 2019). Finally, dying, unlabelled granule cells, located in the superficial granule cell layer and therefore mature or developmentally-born, have been described here and elsewhere (Dayer et al., 2003; Olariu et al., 2005; Cahill et al., 2017).

### The longstanding question of whether the dentate gyrus grows in adulthood

Adult hippocampal neurogenesis has been studied since the 1960s (Altman and Das, 1965) and yet there are only a handful of studies that have addressed the question of whether the DG grows throughout adult life, with no clear consensus on the issue. In the 1980s granule cell growth was first examined by Bayer, who reported that total granule cells increase throughout postnatal life from 1 month to 1 year in rats (Bayer et al., 1982). Bayer estimated that ∼1100 cells are added per day, which aligns with our estimate of ∼800 cells per day. However, Bayer quantified growth beginning *prior to adulthood* (1 month of age). Subsequent work failed to replicate growth over the same interval (Boss et al., 1985) but, given small sample sizes (n=2-4) and a lack of unbiased quantification methods, the issue remained unsettled and, with respect to growth in adulthood, untested.

In the 1990s, the issue was revisited with modern, unbiased stereological counting methods. Interest in hippocampal age-related pathology led to questions about whether age-related cognitive impairments might be explained by neurodegeneration (Rapp and Gallagher, 1996; Rasmussen et al., 1996; Merrill et al., 2003), and whether neurogenesis could provide a source of plasticity in aging (Kempermann et al., 1998).

Some of these studies likely failed to detect growth because the youngest rodents were substantially older (5-6 months) than those used here (2 months)(Rapp and Gallagher, 1996; Kempermann et al., 1998; Merrill et al., 2003). Since the majority of ABNs are added early in adulthood, this would miss a large proportion of the total neurogenic growth (here, 57% of our ABN addition occurred between 2-6 months). In some cases, small sample sizes likely prevented detection. For example, Rasmussen et al. only examined a subset of their behaviorally-tested rats (5 rats/group) (Rasmussen et al., 1996). Reanalyzing their data (extracted with Plot Digitizer) we observe 317,000 more cells at 24 months compared to 3 months. This difference was borderline significant (P=0.046 in our hands) and so growth may have been detected if a larger sample was examined.

Extending these data to other non-human mammals, there is evidence that DG granule cells increase in number in wild populations of bank voles and wood mice (Amrein et al., 2004). The only caveat is that age was determined indirectly (e.g. by tooth wear, evidence of pregnancy, etc) and so there is no way to know precise age. In rhesus monkeys, one study found that granule cells increase in number during infancy, but do not change from 1 to 5-10 years of age (Jabès et al., 2010). However, a study that examined a larger cohort, over a more complete segment of adulthood (5 to 30 years) found significant granule cell addition (Ngwenya et al., 2015).

Human data is limited to a small number of studies reporting that human DG neurons do not increase in number in adulthood (West, 1993; Šimić et al., 1997; Harding et al., 1998; Boldrini et al., 2018). These studies often focussed on age- or neurodegeneration-related cell loss, and therefore many “young” subjects were well into middle age, when neurogenesis may have already declined. However, even in cases where there was a large range of ages, and the youngest subjects were less than 20 years old, there were no trends for an age-related increase in DG neuron numbers (West, 1993; Boldrini et al., 2018). One possibility is that there are higher levels of cell turnover in humans, which completely cancels out neurogenic growth (Spalding et al., 2013). Alternatively, since DG neurogenesis occurs earlier in humans than in rodents, net growth may only be apparent at younger ages (Snyder, 2019).

Translating these findings to humans may be relevant for interpreting hippocampal volume dynamics in clinical situations. For example, hippocampal atrophy is observed in depression (McKinnon et al., 2009) and schizophrenia (Harrison, 2004), and hippocampal growth is seen following electroconvulsive therapy (Nordanskog et al., 2010). It is often speculated that adult neurogenesis could underlie these volume changes though it is unlikely that increased/decreased neuron addition is sufficient to fully explain these volumetric changes. Evidence from animal models indicates that neurogenesis is required neither for stress-related reductions (Schoenfeld et al., 2017) nor electroconvulsive shock-induced increases (Abe et al., 2024), in volume. On the other hand, the total number of DG neurons increases in depressed patients that were treated with antidepressant drugs (Boldrini et al., 2013). Our data suggest that the turnover of DBNs is another factor, in addition to neurogenesis, that could be contributing to volume changes in psychiatric conditions.

### Neurogenesis, neuronal turnover and relevance for long-term memory

The relationship between neuronal turnover and memory persistence is best illustrated in birds, where neuronal turnover in the song system parallels the adoption of new song repertoires (Kirn and Nottebohm, 1993; Cohen et al., 2016). In mammals, although the precise role of the hippocampus in long-term memory remains debated, many theories propose a recurring or permanent contribution of the hippocampus to episodic memory processing and storage (Moscovitch and Gilboa, 2021). If the songbird is an example, net growth of the DG could increase hippocampal storage capacity and allow for the accumulation of episodic memories throughout life. Conversely, with time, many memories become hippocampus-independent, either through forgetting or through transformation via systems consolidation. The selective removal of DBNs may therefore represent one mechanism by which older memory traces are cleared.

An unresolved question is whether causal links exist between the addition of ABNs and the loss of DBNs, as is the case in other systems (Thompson and Brenowitz, 2009; Larson et al., 2014) and cell types (Hughes et al., 2013). Adult neurogenesis is known to promote forgetting, potentially by disrupting existing hippocampal circuits (Akers et al., 2014). Because ABNs appear to compete with established neurons for synaptic space, it is possible that DBNs are actively removed as new neurons integrate into the network (Toni et al., 2007, 2008; McAvoy et al., 2016; Adlaf et al., 2017).

Consistent with this, we found fewer caspase^+^ apoptotic cells in 18-month-old rats, when neurogenesis is low and most ABNs would have fully matured. Also, training in a socially transmitted food preference task both promotes neurogenesis and induces the death of superficially located, putatively developmentally-born granule cells (Olariu et al., 2005). Whether new learning selectively promotes the removal of pre-existing neurons that encode outdated or irrelevant information remains unknown but represents an important direction for future studies of hippocampal plasticity and memory.

## Notes

### Competing Interest Statement

The authors have declared no competing interest.

### Funding

- Natural Sciences and Engineering Research Council
- Canada Foundation for Innovation
- Michael Smith Health Research BC

